# Cortical activity at different time scales: high-pass filtering separates motor planning and execution

**DOI:** 10.1101/857300

**Authors:** David Eriksson, Mona Heiland, Artur Schneider, Ilka Diester

**Affiliations:** Optophysiology, University of Freiburg, Faculty of Biology, 79104 Freiburg, Germany; Optophysiology, University of Freiburg, Faculty of Biology, BrainLinks-BrainTools, Bernstein Center Freiburg, 79104 Freiburg, Germany

**Author notes:** Department of Physiology and Medical Physics, Royal College of Surgeons in Ireland | RCSI, 123 St. Stephen’s Green, Dublin 2, Ireland.

## Abstract

The smooth conduction of movements requires simultaneous motor planning and execution according to internal goals. So far it is not known how such movement plans can be modified without being distorted by ongoing movements. Previous studies have isolated planning and execution related neuronal activity by separating behavioral planning and movement periods in time by sensory cues^1–7^. Here, we introduced two novel tasks in which motor planning developed intrinsically. We separated this continuous self-paced motor planning statistically from motor execution by experimentally minimizing the repetitiveness of the movements. Thereby, we found that in the rat sensorimotor cortex, neuronal motor planning processes evolved with slower dynamics than movement related responses both on a sorted unit and population level. The fast evolving neuronal activity preceded skilled forelimb movements while it coincided with movements in a locomotor task. We captured this fast evolving movement related activity via a high-pass filter approach and confirmed the results with optogenetic stimulations. As biological mechanism underlying such a high pass filtering we suggest neuronal adaption. The differences in dynamics combined with a high pass filtering mechanism represents a simple principle for concurrent motor planning and execution in which planning will result in relatively slow dynamics that will not produce movements.

## Main Text

In smooth movement sequences, a continuum from motor planning over motor execution to sensory integration can be defined, according to the temporal lag between neuronal activity and behavior. Here, we consider neuronal activity with a temporal lag to the behavior in the order of a previously suggested range of less than 100 ms^8,9^ as being related to motor execution. We refer to neuronal activity with larger temporal lags to the behavior as motor planning or sensory integration, depending on whether the neuronal activity occurred before or after the movement. This lag based interpretation of neuronal processes is hampered by behavioral correlations. If two behavioral processes are correlated (e.g. because they always occur in the same sequence), neuronal activities appear to be correlated to both even if a causal relationship only exists for one of the behavioral processes. To reduce this temporal bleeding, we aimed to minimize correlations by encouraging animals to conduct movements with minimal reoccurrence of individual movement sequences. In the locomotor task, rats moved unconstrained in a box while searching for pseudo-randomly placed water drops on a floor mesh (**Fig. 1A**). In the joystick task, rats were trained to move a joystick with their right front paw while minimizing revisiting previously visited positions (**Fig. 1B**). Thus, rats had to internally develop movement plans to optimize the number of rewards. In both tasks, movements were not repetitive as indicated by the narrow temporal behavioral autocorrelations of the movement velocities (see data boxes in **Fig. 1C and D**). A repetitive movement or prolonged behavioral state would cause an autocorrelation with multiple peaks (see illustration in **Fig. 1C)** or one broader peak due to experimentally induced delay periods which can lead to an extended neuronal activity often interpreted as motor planning (see illustration in **Fig. 1D**), respectively.

**Figure 1.**
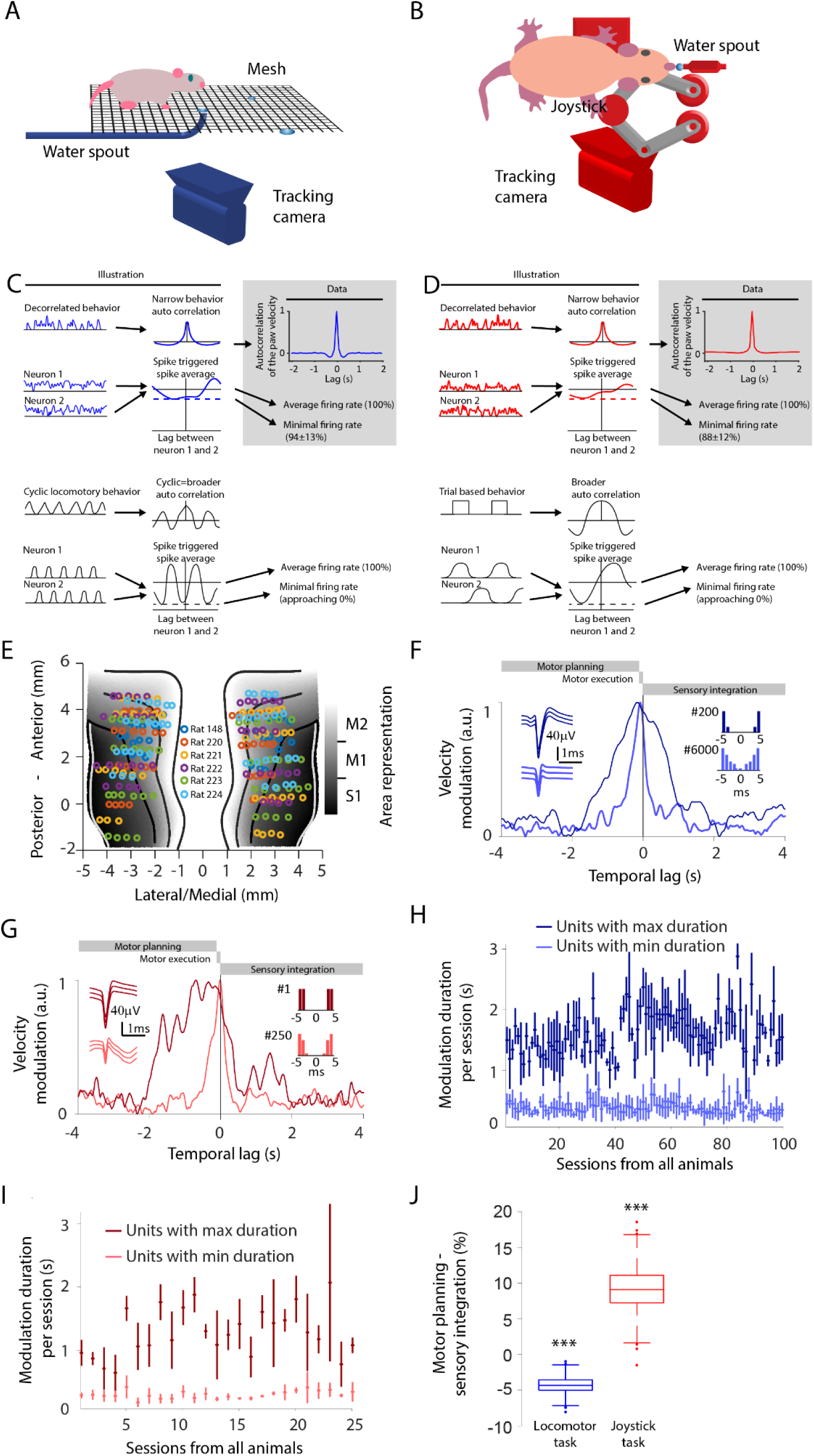
Studying neuronal dynamics with minimally repetitive behavior. A: Setup of locomotor task. B: Setup of joystick task. C: Illustration of the difference between decorrelated and repetitive behavior in terms of the behavioral autocorrelation and neuronal cross correlation for the locomotor task. The behavioral autocorrelation is broader for a repetitive locomotion (bottom panels) than for a decorrelated behavior (top panels). The minimal value (dashed line) of the neuronal cross correlation is low if there are lags for which the two neurons do not spike (indicating correlated firing) and it is high if the two neurons fire at different lags (indicating de-correlated firing) (illustration in left panel). Autocorrelation for the velocity of the right front paw during the locomotor task (gray data panel). D: Same outline as in C but for the joystick task. A repeating trial structure causes correlations between different trial periods. This in turn may increase the width of the behavioral autocorrelation as well as the correlation between neurons. E: Electrode locations on the sensorimotor cortex for respective animal. F: Velocity modulation of the instantaneous firing rate for 2 example units with action potential waveforms (left inset) and interspike interval histogram (right insets) in the locomotor task. Neuronal firing rates modulated by future or past paw movement velocities are assigned to negative temporal lags (referring to planning) or to positive temporal lags (referring to sensory integration), respectively. Lags between 0 and 100ms are considered to be related to motor execution. The dark-blue unit has a broad velocity modulation, whereas the light-blue unit is temporally very precise. Both units originate from the same recording session. G: Same outline as in F but for two different units in the joystick task. The dark-red unit has a broad velocity modulation, while the light-red unit is temporally precise. H: The unit with the minimal (bright-blue) and maximal (dark-blue) duration of the velocity modulation for each locomotor session. The error bars denote the standard deviation of bootstrapped durations. I: Same outline as in H but for the joystick task. Light-red and dark-red corresponds to units with minimal and maximal modulation duration respectively. J: The summed velocity modulation for motor planning-related activity (negative lags from -1.1 to -0.1 s) minus the summed velocity modulation for sensory integration related activity (positive lags from 0 to 1 s). Significances are indicated according to: *** p < 0.001.

To study the neuronal underpinnings of decorrelated movements, we trained six Long-Evans rats in the locomotor task. Five of these animals were also trained in the joystick task. To record neuronal activity, electrodes were placed bilaterally in the sensorimotor cortex (42 electrodes per animal, **Fig. 1E**). We targeted the output layer V by implanting the electrodes at a depth of 1.2 mm^10,11^. In total we recorded 5400 single units (SU) and 6876 multi units (MU) over 100 sessions for the locomotor task (**Supplementary Table 1**) and 1217 SU and 1659 MU over 25 sessions for the joystick task (**Supplementary Table 2**). We refer to SU and MU collectively as sorted units. For repetitive behavior, neurons may fire at a specific lag relative to each other, rendering some lags less represented than others. This causes the firing rate for some lags to be fundamentally lower than the average firing rate (see dashed lines in **Fig. 1C and D**). Here, the neuronal activity was characterized by a decorrelated pair-wise spiking, i.e. pairs of neurons fired independently of each other such that all lags were represented equally, and the firing rate at a certain lag was close to the average firing rate. The firing rate of one neuron relative to another neuron at the least represented lag was 94 ± 13% and 88 ± 12% of the average firing rate in the locomotor and joystick task, respectively (**Fig. 1C and D**, see methods) indicating a decorrelated neuronal activity. Because both the behavioral and the neuronal activity were decorrelated with respect to time, the temporal bleeding was minimized and the temporal precision of the estimated functional relation between movement and neuronal activity was optimized. To quantify the temporal precision, we calculated the range of temporal lags for which a given sorted unit was modulated by the paw velocity (**Fig. 1F and G**). We refer to this modulation across lags as velocity modulation and the duration for which the velocity modulation exceeded 80% of the peak modulation we refer to as the modulation duration. We observed units with both long modulation durations (locomotor task: 1.6 ± 0.37s, joystick task: 1.2 ± 0.37s) and short modulation durations (locomotor task: 0.36 ± 0.09s, joystick task: 0.27 ± 0.06s) within the same session (**Fig. 1H and I**). This demonstrates that our approach minimized behavioral bleeding to the extent which allowed separating long processes, like motor planning and sensory integration, from shorter processes like motor execution. Finally, this behavioral approach enabled us to quantify the relative strength of motor planning and sensory integration by taking the normalized difference of the velocity modulation for negative and positive temporal lags. In line with previous lesion and inactivation approaches^12–15^, the relative contribution of the motor planning related activity was larger for the joystick task (9.3 ± 2.8%, mean ± SEM, p=0.0007, two-tailed t-tests) whereas the sensory integration related activity was larger in the locomotor task (−4.3 ± 1%, mean ± SEM, p<0.0001, two-tailed t-tests, **Fig. 1J**). Thus, our approach based on minimal repetitive movements complements previous studies with a temporally refined neuronal activity based assay of the gradient from motor planning and execution to sensory integration for skilled and locomotor behavior.

### Varying neuronal modulation durations

Motor execution can be generated by sequentially activated sets of neurons, or similarly, a sensory event may traverse through the network. A temporal recruitment of neurons has been described for attractor networks^16,17^. Those studies focused on a special case in which each neuron was activated for a constant duration (**Fig. 2A, upper panel**). Alternatively, the modulation duration may increase with larger temporal lags relative to the movement (**Fig. 2A, lower panel**). Here we defined the temporal lag based on the peak of the velocity modulation (see methods). In accordance with the second hypothesis, the modulation duration increased significantly with increasing temporal lags for both locomotor and joystick tasks (ANOVA, locomotor task, p<0.0001, ANOVA joystick task, p<0.0001, **Fig. 2B and C**). This suggests that putative motor execution represented by units with shorter temporal lags occurred with faster neural dynamics than motor planning and sensory integration.

**Figure 2.**
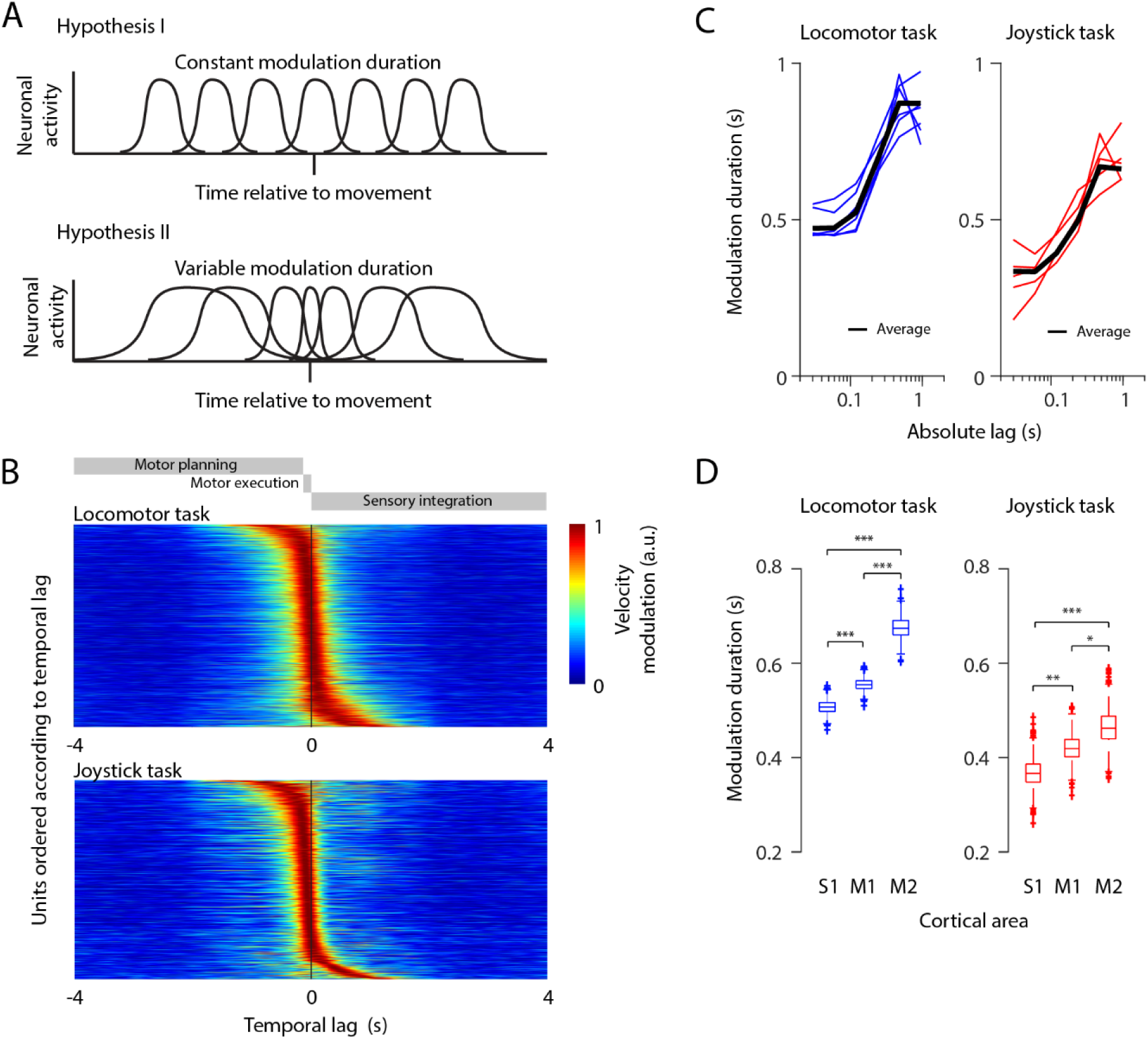
The duration of velocity modulation of individual units depends on the temporal lag to behavior and on the cortical area. A: Two hypotheses of sequential neuronal activity relative to movement. The duration of the neuronal activity can be constant across lags (upper panel) or different across lags (lower panel). B: Units sorted according to the lag of their maximum velocity modulation in the locomotor task (top), and for the joystick task (bottom). C: Relation between average modulation duration and temporal lag (black line), across the different animals (colored lines) for the locomotor (left) and joystick task (right). D: Duration of the velocity modulation for each cortical area for the locomotor task (left) and the joystick task (right). Significances are indicated according to: * p < 0.05, ** p < 0.01, and *** p < 0.001.

### Integration timing of cortical areas

If motor planning and sensory integration is associated with longer modulation durations, it is conceivable that a higher brain area, such as secondary motor cortex (M2, putatively functionally similar to premotor cortex in primates^18,19^) contains neurons with longer modulation durations than primary motor cortex (M1). To test this, we mapped the electrode locations on to the non-linear gradient spanning M2, M1, and primary somatosensory cortex (S1) (**Fig. 1E**). Indeed, neurons in higher areas (i.e., M2) had a significantly longer modulation duration than neurons in lower areas (i.e., M1 and S1, **Fig. 2D**). This was true for both the locomotor and the joystick task (ANOVA, locomotor task: p<0.0001, joystick task: p<0.0001). On average, neurons in S1, M1, and M2 had a modulation duration of 507±14, 555±13 and 676±22 ms during locomotor and 369 ± 29, 423 ± 27, and 469 ± 38 ms (mean ± SEM) during the joystick task.

### Population activity destabilizes during movement

Next, we examined whether the differences between the two tasks regarding the modulation duration of individual units also generalized to the neuronal population. To this end, we correlated the population activity, including all sorted units at any two time points. We refer to this correlation as population correlation. The population correlation will typically decay with increasing temporal distance between the two time points. This population correlation decay is a measure for the stability of the population activity, i.e. how slow (stable over time) or fast (instable over time) the population activity changes. To compare the stability of the population activity during movement and behavioral quiescence, we defined trials between the time point of lowest paw velocity (peri-trial time of - 1 second) which we refer to as premovement and the time point of highest paw velocity (peri-trial time +1 second) which we refer to as movement (see methods, **Fig. 3A and B**). While the population correlation followed a similar motive with a less confined diagonal during premovement and a more confined diagonal during movement, robust bands of low correlation during movement execution only occurred in the joystick task, but not in the locomotor task, thus revealing a qualitatively different correlation structure (**Fig. 3C and D**). These bands of low correlation are a sign of a quick decay of the population correlation, indicating that the population activity changed rapidly during motor execution. To quantify how fast the population correlation decayed, we fit an exponential function to the decay of the population correlation. During periods of movements, population correlations decayed significantly faster than the median time constant in the joystick task (−176±59 ms, mean ± SEM, n=5, p=0.043, two-tailed t-tests) but not in the locomotor task (−18±27 ms, mean ± SEM, n=6, p=0.54, two-tailed t-tests, **Fig. 3E and F**). In line with the strong decrease in time constant in the joystick task during movements (**Fig. 3G**), the time constant during joystick movements was lowest (203±88 ms, mean ± SEM, n=5, **Fig. 3H**) indicating an unstable population activity. In contrast, the time constant was largest (i.e. the population activity was stable) during joystick premovement periods which putatively involves motor planning (761±375 ms, mean ± SEM, n=5, **Fig. 3H**). The difference in stability of the population activity cannot be explained by behavioral differences across the two tasks (summarized in Supplementary Note 1). To summarize, this suggests that premovement periods (putatively involving motor planning) were associated with stable population activity with slow changes in the neuronal activity whereas movements (referring to motor execution) were associated with unstable population activity with fast changes in the neuronal activity.

**Figure 3.**
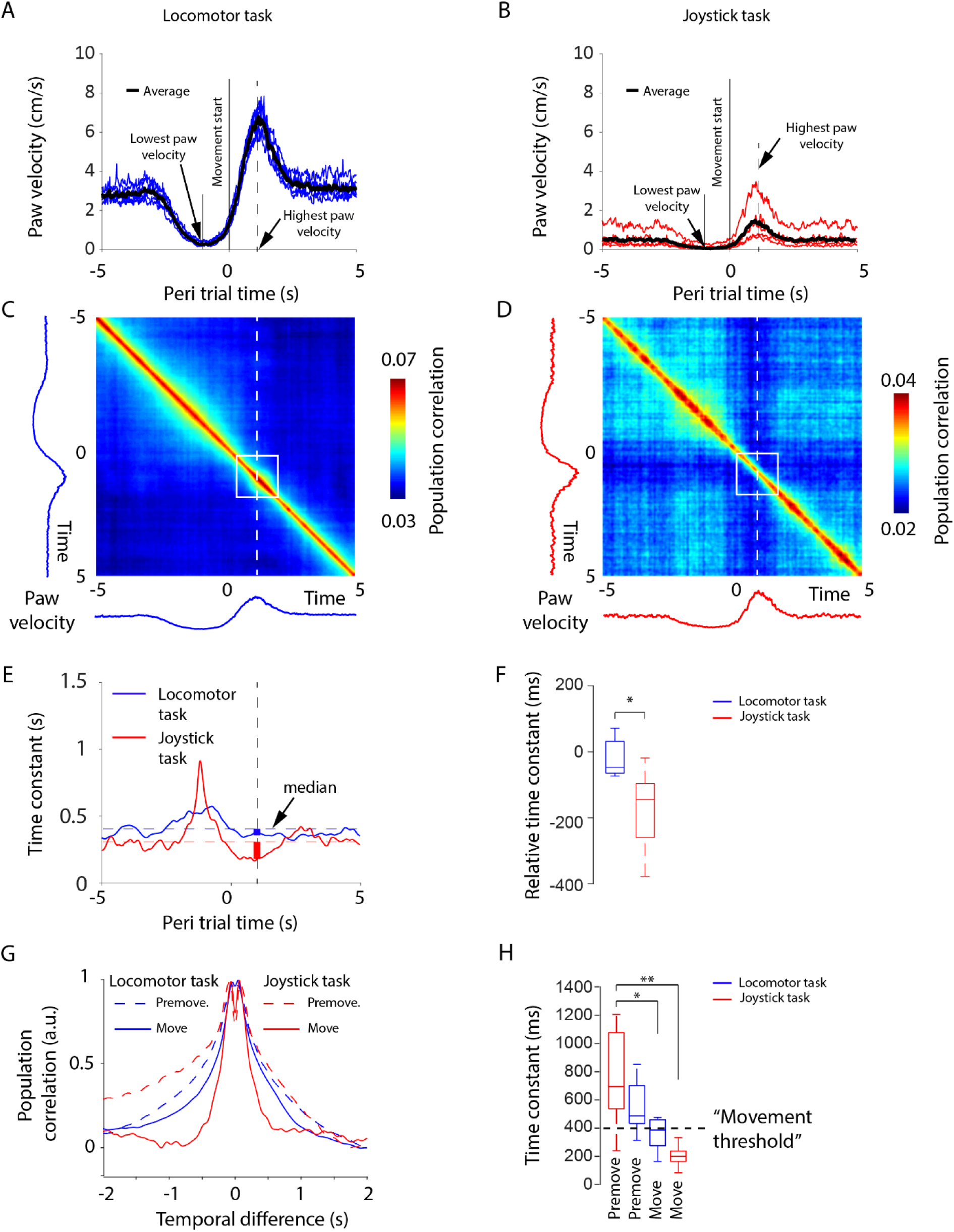
The population activity changes faster during motor execution in the joystick task as compared to the locomotor task. A: Average paw velocity across behavioral trials for the locomotor task (see methods for the trial definition). Blue lines denote data from individual animals and the black line denotes the average across all animals. B: Same outline as in A but for the joystick task. C: Average pairwise correlations of population vectors across all animals for the locomotor task. D: Same outline as in C but for the joystick task. E: The time constant of the decay in the population correlation across the trial in the locomotor task (blue) and the joystick task (red). The median relative time constants are included as dotted lines. F: The decrease in time constant during movement in relation to the median time constant across the trial for locomotor and joystick task. G: Decay in population correlation at the premovement time point (at -1 second during lowest paw velocities) and at the movement time point (at 1 second during highest paw velocities). H: Time constants for the locomotor and joystick task for the premovement and movement time points based on the curves in G. Significances are indicated according to: * p < 0.05, ** p < 0.01.

### Fast changes in neuronal activity precede movement

Unstable population activities associated with motor execution could be captured in the high frequency range. Thus, a high-pass filtered neuronal activity should be tightly correlated to movement execution (**Fig. 4A**). Paw velocities provide a general measure of movement magnitude independent of specific types of movements. To allow a comparison of the discretized and typically low frequency spike trains of sensorimotor cortex with the continuous paw movements, we reconstructed the continuous subthreshold activity with a resolution of 10 ms from the spiking activity^20^ (**Fig. 4B**). This allows the detection of neuronal activity changes which are faster than those signaled by low frequency spiking events. Fast changing activities typically preceded large paw velocities (**Fig. 4C**). To quantify this relation, we calculated the Pearson correlation coefficient between paw velocity and the rectified high pass filtered neuronal activity (averaged across neurons, cut off frequency 1.1 Hz) (**Fig. 4D**, upper panels). The cut off frequency of 1.1 Hz was motivated by the maximum correlation at this frequency (**Fig. 4E**) The high pass filtering was contrasted against corresponding calculations for the low pass filtered neuronal activity (**Fig 4D**, lower panels). The correlation was generally higher for the high pass filtered neuronal activity than for the low pass filtered neuronal activity both for the locomotor task (low pass: 0.059±0.029 vs. high pass: 0.17±0.025, mean ± SEM, n=6, p=0.0163, two-tailed t-tests, **Fig 4D, E and F**), and for the joystick task (low pass: 0.1±0.025 vs. high pass: 0.20±0.013, mean ± SEM, n=5, p=0.0091, two-tailed t-tests). For the joystick task, the correlation reached its maximum at a small negative lag between high pass filtered neuronal activity and movement, which falls in the range of movement execution (−94 ± 20 ms, mean ± SEM, p = 0.01, n = 5, two-tailed t-tests, **Fig 4D and G**), whereas the peak for the locomotor task did not significantly precede the movement for any frequency band (120 ± 110 ms, mean ± SEM, p = 0.33, n = 6, two-tailed t-tests). The lag of the peak of the correlation was significantly shifted to positive values corresponding to sensory integration for the low-pass filtered neuronal activity in the locomotor task (770±117 ms, mean ± SEM, p = 0.0013, n = 6, two-tailed t-tests), and to negative values corresponding to motor planning components in the joystick task (−234±36 ms, mean ± SEM, p = 0.0029, n = 5, two-tailed t-tests). Thus the frequency of the neuronal activity changes separated planning and sensory integration from motor execution.

**Figure 4.**
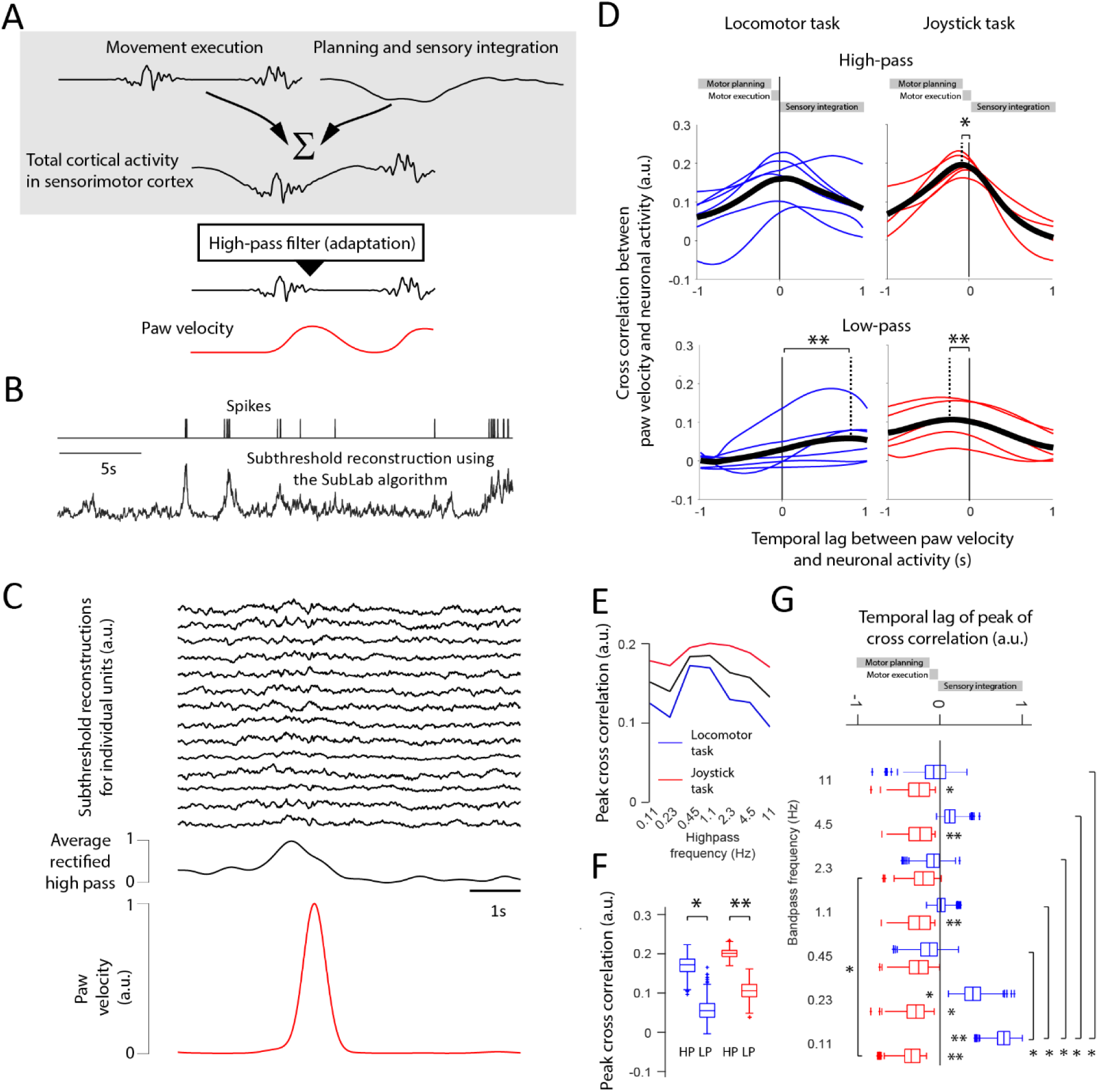
High-pass filtered neural activity is correlated to paw velocities. A: Schematic illustration of how a high-frequency neuronal activity can be superimposed on a low-frequency neuronal activity and yet be separable. B: To be able to study how fast neuronal activities change, we reconstructed the subthreshold activity of the sorted units. C: Reconstruction of neuronal activities from 14 randomly selected units during the joystick task (top). An increase in the average rectified high pass filtered neuronal activity (black trace, middle row) typically precedes higher paw velocity (red trace). D: The Pearson correlation coefficient for different lags between the rectified high-pass filtered neuronal activity and the paw velocity during the locomotor task (upper-left), and the joystick task (upper right), and for the low-pass filtered neuronal activity and the behavior during the locomotor task (lower left), and the joystick task (lower right). The comparison between cross-correlation values at time point zero and the time point of maximal cross-correlation reveals significant changes with different temporal lags. E: The peak correlation between the paw velocity and the rectified high pass filtered neuronal activity for seven cut off frequencies for the joystick task (red), the locomotor task (blue) and their average (black). F: Peak Pearson correlation coefficients for panel D. G: Temporal lags of the peak Pearson correlation coefficient (across temporal lags) for band-pass filtered neuronal activity. Significances are indicated according to: * p < 0.05, ** p < 0.01.

Since previous work has separated planning and motor execution in terms of a population code^3^ we asked if the frequency code could predict this population coding. The tuning of the paw velocity (**Fig. S3A** and **B**) was correlated to the change in population code when traversing from motor planning towards motor execution (**Fig. S3C**). Since the correlation between the population velocity tuning at planning lags (−1000 to -200 ms) and at execution lags (−40 ms) was minimal we regarded the population velocity tuning at -40 ms to represent the output potent space. The space orthogonal to this was regarded as the output null space. Indeed, the part of the trajectory that corresponded to high frequency coding was typically associated with high paw velocities and represented the output potent space. In contrast, the part of the trajectory that corresponded to low frequency coding was typically characterized by low paw velocities and lied in the output null space on a single sessions level (**Fig. 5A. Fig. S3D**) as well as in an across session average (high frequency coding: p<0.00001, Bonferroni corrected, n=929, low frequency coding: p< 0.00001, Bonferroni corrected, n=579, **Fig. 5B**). Thus the frequency coding can predict population code indicating that frequency coding may be an integral part of movement control which might represent the underlying biological explanation of the population code theory.

**Figure 5.**
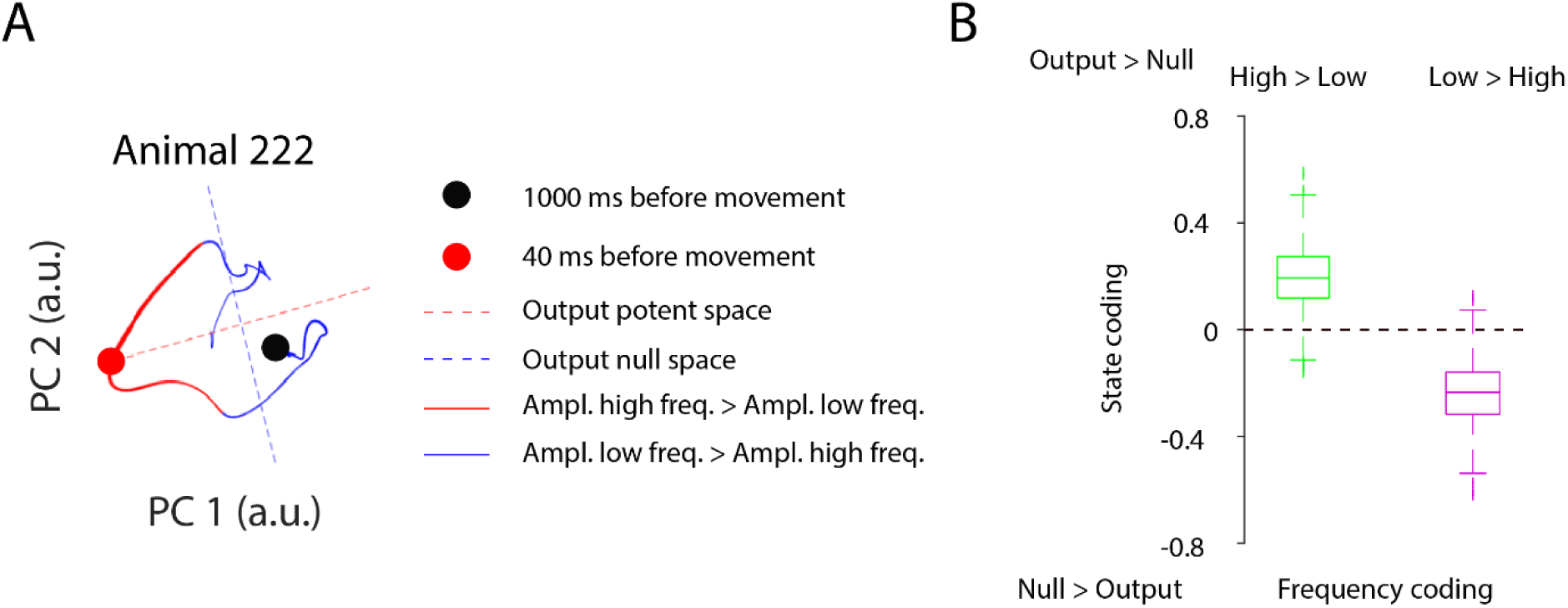
Ratio of high to low frequency changes of neuronal data predicts state spaces. A: Example sessions of dimension reduced population coding for animal 222. The trajectory is divided into paths for which the high frequency (>1.1Hz) had a larger amplitude than the low frequency (<1.1Hz) (red), and into segments for which the high frequency had a smaller amplitude than the low frequency (blue). Output null (blue) and output potent spaces (red) are indicated by dashed lines. The thickness of the trajectory indicates the averaged paw velocity. Thicker lines refer to higher velocities. B: Quantification of the results in panel A for all animals. The average state coding was calculated for preferentially high frequency and low frequency coding. A negative or positive state coding value refers to a dominant null space or output potent space, respectively. The frequency coding is defined by either a dominance of high or low frequencies of neuronal changes. Dominant high frequency changes are associated with the output potent space, whereas dominant low frequency changes were correlated with the null space.

### High frequency optogenetic stimulation evokes movements

Next we optogenetically induced brain activity in the primary motor cortex at different frequencies to examine whether a certain frequency was more prone to evoke paw movements. We tested five different frequencies 0.1, 0.3, 1, 3, and 10 Hz. To minimize the effect of harmonics, the light was varied according to a sinusoidal. To test whether slow oscillations induced a depolarization block, we measured extracellular activity in close proximity to the optical fiber (**Fig. 6A** and **B**). All stimulation frequencies resulted in a strong increase of neuronal firing. However, in line with our high frequency hypothesis, only 3 and 10 Hz resulted in an overt cyclic paw movement (**Fig. 6C** and **D**, supplementary movie 1). This movement threshold between 1 and 3 Hz is in line with the fact that movement generation was associated with a time constant of 400 ms (**Fig. 3H**) which in turn corresponds to a 2.5 Hz, and since the peak in the correlation between paw velocity and neuronal activity occurred at 1.1 Hz (**Fig. 4E**).

**Figure 6.**
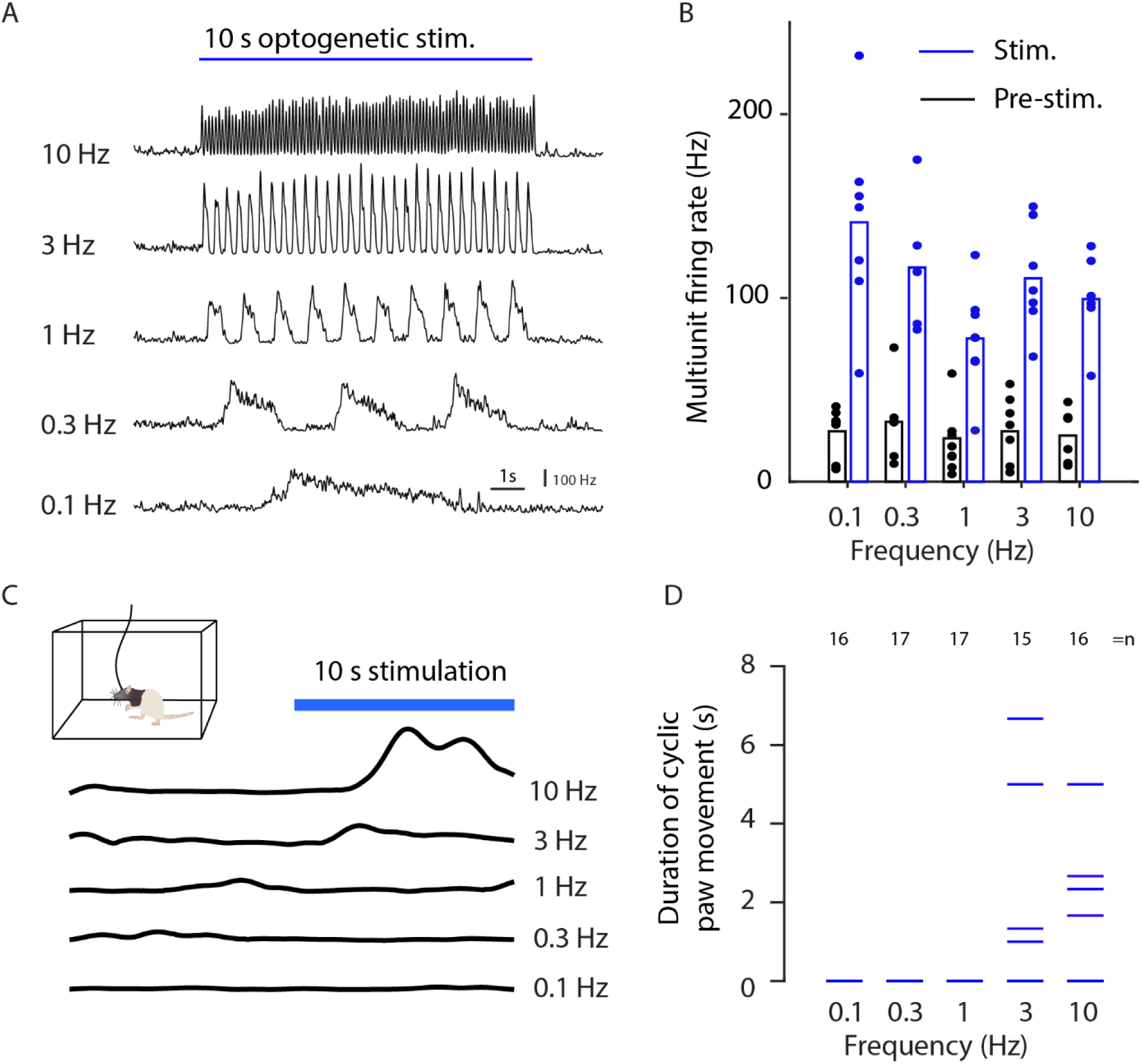
Optogenetic induction of movement. A: Simultaneous extracellular recordings and optogenetic stimulation at five different frequencies averaged across two animals. The instantaneous firing rate was estimated by threshold crossings in 10 milliseconds bins for the stimulation period of 10 seconds. B: Average firing rate before (black) and during stimulation (blue) for different trials (dots) from two animals. The bars indicates average firing rate. C: Optogenetic stimulation in a plexiglas arena surrounded by 7 cameras for behavioral tracking^31^. The behavior was quantified as the amplitude of the velocity for respective stimulation frequencies (black lines). D: Behavioral quantification for different stimulation frequencies. Each line correspond to the duration of cyclic movements of the paw in a stimulation trial. Optogenetic stimulation did not evoke any cyclic movements in the trials for frequencies 0.1 (16 trials), 0.3 (17 trials) and 1 Hz (17 trials). Only optogenetic stimulation applied with 3 or more Hz induced overt cyclic movements.

## Discussion

Based on two tasks that encouraged animals to conduct minimally repetitive movements, we found that fast changes in neuronal activity were related to motor execution. These fast changes in neuronal activity were more pronounced during the joystick task than during the locomotor task. Furthermore, higher frequencies in the neuronal activity preceded the movement by 100 ms in the joystick task, whereas it coincided with the movement for the locomotor task. This is in line with the fact that the locomotor task required no training (**Fig S1A**) and that locomotion may be dominated by an efference copy signal in the neuronal activity^15,21^. In contrast, the joystick task required training (**Fig S1B**) and lesioning and inactivation studies have shown that skilled movements are more dependent on the motor cortex^12–14^. Here we showed that lower frequencies were decoupled from movement suggesting that they were more related to motor planning and sensory integration. Movement decoupled activity avoided the high frequencies underlining the general role of the fast changing neuronal activities in motor cortex for movement execution. Such a fast changing population activity can be extracted by a classic high pass filter. Fast changes refer to e.g. changes from a high firing rate to a low firing rate, or vice versa. Adaptation mechanisms^22–26^ at any stage between the cortex and the muscles could serve as the biological equivalent of such a high pass filter (see Supplementary Note 2). As a direct test of our frequency hypothesis optogenetic stimulation only resulted in movements when applied at sufficiently high frequencies.

The here proposed frequency based separation of motor planning and execution can be integrated into conceptual frame works of motor control. According to the concept of dynamical systems, e.g. the null-space theory^3^, the frequency based separation of motor planning and execution would allow both processes to work in parallel. So far the null-space theory was tested with trial structures with temporally separated planning and execution periods^6^ or with sensory driven motor execution^27^. For intrinsically planned continuous movements, our results suggest that two independent population state spaces can be generated in the frequency domain, one based on high and one on low frequencies. The concept of separate neuronal populations for motor execution and motor planning (e.g. by genetically or projection defined neurons^1,2^) assumes a complete separation of the signals. However, genetically defined spinal cord projecting neurons have been shown to not only encode motor execution but also motor planning^2,7^. Our proposed high-pass filtering mechanism could be a way to expose the motor execution component by decreasing the planning component. Therefore, the separation of the processes by means of slow and fast dynamics could facilitate simultaneous parallel motor planning and execution within the same neuron, be it in the conceptual frame work of dynamical systems or based on identified neuronal subtypes.

The separation of motor planning and execution by means of different frequencies of neuronal activity requires that motor planning evolves relatively slowly. This prerequisite is reasonable, as planning and decision making rely on accumulating internal or external evidence^28–30^. Thus, motor planning-related neuronal activity changes slowly and hence can be stopped from percolating to the muscles by a high pass filtering mechanism based on neuronal adaptation. Thus, our proposed mechanisms is able to explain in a very simple manner the simultaneous implementation of intrinsic motor planning and execution.

## Supporting information

Supplementary Infomration

## Methods

### Animals

All animal procedures were approved by the Regierungspräsidium Freiburg, Germany. In this study we used six male Long Evans rats (400 g, Janvier) which were implanted at the age of eight weeks and recorded up to four months after the implantation. Three to four animals were pair-housed in type 4 cages (1500U, IVC typ4, Tecniplast, Hohenpeißenberg, Germany) before implantation and the animals were single housed after the implantation in type 3 cages (1291H, IVC typ4, Tecniplast, Hohenpeißenberg, Germany) under a 12 h light dark cycle (dark period from 8 a.m. to 8 p.m., time span of training and experiments). Prior to the first behavioral training, no behavioral tests were conducted, no drugs were applied and food (standard lab chow) and water were provided *ad libitum*. During the course of the experiment, the animals were maintained with free access to food but water supply was restricted. Rats were kept at > 80 % body weight as measured prior to water restriction. For 2 days per week, free access to water was ensured.

### Animal surgery

Animals were initially anesthetized with isoflurane inhalation followed by intra-peritoneal injection of 75 mg/kg Ketamine (Medistar, Holzwickede, Germany) and 50 Μg/kg Medetomidin (Orion Pharma, Espoo, Finland). The animals were then put into a transportation container covered with an opaque cloth to facilitate the anesthesia. Once the animals were anesthetized, they were positioned in a stereotaxic frame (David Kopf Instruments, Tujunga, CA, USA) and their body temperature was kept at 36 °C using a rectal thermometer and a heated blanket (FHC, Bowdoin, USA). The anesthesia of the animals was maintained with approximately 2% isoflurane and 0.5 l/min O2. For pre-surgery analgesia, we subcutaneously (s.c.) administered 0.05 mg/kg Buprenorphine (Selectavet Dr. Otto Fischer GmbH, Weyarn/Holzolling, Germany). Every other hour, the animals received a s.c. injection of 5 mL isotonic saline. Moisturizing ointment was applied to the eyes to prevent them from drying out (Bepanthen, Bayer HealthCare, Leverkusen, Germany). The skin was disinfected with Braunol (B. Braun Melsungen AG, Melsungen, Germany) and Kodan (Schülke, Norderstedt, Germany). To perform the craniotomy, the skin on the head was opened along a 2 cm long incision using a scalpel. The exposed bone was cleaned using a 3% peroxide solution. Self-tapping skull screws (J.I. Morris Company, Southbridge, MA, USA) for reference for extracellular recordings were placed over cerebellum. Craniotomies were drilled bilaterally extending from -2 to +5 mm in the anterior posterior direction and from +1 to +4 mm in the lateral medial direction relative to Bregma. 22 tungsten electrodes (200 to 600 kOhm impedance, polyimide insulation, WHS Sondermetalle, Grünsfeld, Germany) were implanted at a depth of 1.2 mm in each hemisphere. Electrodes were implanted according to the area borders given by the online brain atlas from Matt Gaidica^31^ (**Fig. 1E**). Three rows of 6 electrodes each, oriented in the medial-lateral direction, were implanted in the anterior-posterior direction. The fourth and last row consisted of 4 electrodes, oriented in the medial-lateral direction (see **Fig. 1E**). Occasionally, we had to cut some electrode wires, in order to not destroy blood vessels at the implantation site (e.g., rat 221, left hemisphere, last electrode row). Kwik-Cast (WPI, Sarasota, FL, USA) was used to protect the brain from the dental cement applied in the final step. Before, Mill-Max connectors (Mill-Max, Oyster Bay, USA) from each hemisphere were glued together to form a 4 × 13 pin connection matrix. The last and first four pins were connected to the two skull screws over cerebellum to serve as reference and ground. Finally, the assembly was fixed using dental cement (Paladur, Kulzer GmbH, Hanau, Germany).

### Behavioral tasks

Animals were encouraged to move with as little repetition as possible. In the locomotor task, two servo motors positioned a waterspout at different locations within an arena of 30×40 cm. Every 10 to 30 s a valve ejected a drop of water, which remained in the mesh until the rats consumed it. To prevent the rats from following the movements of the waterspout, we introduced dummy moves: First the waterspout was doing a dummy move without giving water. One second later it did move to a new position where it let out a water drop. The third and last move was again a dummy move. Even for an experienced animal, this procedure resulted in multiple water drops distributed across the mesh at any given time point. The fact that the rats did not collect all water drops indicates that the animals could not predict where the water was let out and had to actively search for it. This task required minimal training as indicated by the stable paw velocities over all sessions. Thus, we used all sessions for data analysis (**Supplementary Fig. 1A**).

In the joystick task, the animals had to learn to grab a joystick-like manipulator as a first step. The manipulator was based on a manipulandum for rodents^32^. Instead of having to reach out for the joystick, the joystick was placed right below the right front paw. The naïve rats typically explored the arena in which the joystick was placed. As the animals placed the paw by chance on the joystick, the joystick vibrated and a liquid reward was given as long as three requirements were met: (1) The rats had to keep holding the joystick with the right front paw which we controlled for via force sensors on the joystick. (2) The left front paw had to be placed on a force sensor plate, which was placed to the left of the joystick. (3) The rats’ head had to cross an infrared sensor. This ensured that the animals had to learn to use their right front paw to manipulate the joystick rather than the left paw or the mouth. The vibration of the joystick was implemented by clamping the current of the two motors according to two independent Gaussian processes and served two purposes: (1) it made the animals aware of the joystick. (2) The vibration of the joystick increased in amplitude during the course of 10 s (the maximum vibration amplitude resulted in an average acceleration of 1.5m/s^2^) such that, unless the animals held the joystick firmly, it would lose the grip and thus not receive rewards. Together, these measures resulted in an automatic training by which the rats learned to hold the joystick during the maximum vibration amplitude within 10 sessions. Once the rats had developed a firm grip of the joystick, the motors were turned off and the rats received a reward when they actively moved the joystick. Moreover, the rats only received rewards when they moved in a direction or to a position which had not been visited recently (see below). The joystick could be moved within an arena of 40×40 mm. This arena was divided into 5×5 bins and the direction of movement was divided into 8 bins. For each bin we stored the amount of remaining reward. Whenever the rats visited one bin, the amount of remaining reward, r, in that bin was decreased to r-Δr. The amount of reward that was decreased, Δr, was distributed among all other bins. Thus, if the rats preferred one bin, the reward within that bin disappeared completely after 20 seconds. It took up to 15 sessions for the animals to start to move the joystick non-repetitively (**Supplementary Fig. 1B**). Before the rats started to move randomly, they typically tried to pull the joystick only in one direction (typically towards the rat). This resulted in minimal overall movements since the joystick was stopped by the edges of the arena (the 40×40 mm arena). Only when they realized that they could move in all different directions, the amount of total movement increased. For data analyses, we used data from sessions 15 to 35.

### Quantifying behavior

Since the rats had to take a defined pose in the joystick task, we could relate the joystick position and movement to the egocentric coordinates of the rat. To enable a comparison of the locomotor task and the joystick task, it was necessary to quantify the behavioral variables in a similar way. To achieve an egocentric tracking in the locomotor task, we tracked the paws, head, chest, and belly of the animals. By using the head, chest, and belly coordinates, we aligned the movements of the right front paw to egocentric coordinates. Those body parts were tracked by painting them in different colors. The head of the rat did not have to be painted because of the black hood of Long Evans rats. To ensure that all body parts could be tracked, the cameras were placed below the arena. Two to four cameras (Stingray, F033C IRF CSM, Allied Vision Technologies) were used in the locomotor task. The noise of tracking was estimated to 0.79 cm/s (estimated when the paw was standing still on the mesh) and was subtracted from the paw velocity estimates.

### Data acquisition and preprocessing of extracellular recordings

Extracellular signals were bandpass filtered, amplified and digitized using the INTAN (Intan Technologies, Los Angeles, California) head stage attached to the Mill-Max matrix connector at the head of the animals. To maximize comfort for the animals, we stripped the ultrathin INTAN cable and suspended it with a 1.5 m long ultralight spring with a 1.5 mm diameter. The long recording cable allowed the rats to move between the locomotor task and the joystick task without having to be disconnected and re-connected. The rats could either begin with the locomotor task and after 30 min a door was opened allowing the rats to walk into the joystick arena for 40 to 90 minutes, or the rats were in the joystick arena for the entire session. In case of a dual task session, we always began with the locomotor task, because the color markers used for the locomotor tracking faded over time.

The extracellular recordings were sampled at 30 kHz and were de-noised offline. First, 50 Hz and the corresponding harmonics were removed using a 20 ms template estimation. The activity across all channels was demeaned using a median filter. Spike sorting was conducted on high-pass filtered data with a cut off frequency of 300 Hz. Spike snippets were extracted from peak aligned events that crossed a threshold of four times the standard deviation. Only spikes with a negative peak were taken into account. The spike window was -0.5 to 2 ms around the peak amplitude (resulting in 76 values for each spike). To minimize the risk that a sorted unit was a combination of multiple neurons, we applied a conservative threshold for the cluster size. To this end we used a cluster size that was dictated by the noise level half a millisecond before the minimum of the spike. Given the typical refractory period of neurons, this noise estimate excluded variability caused by this unit and was therefore a direct measure of the cluster size of this particular unit. Since our electrodes typically had a spacing between 300 and 1000 µm, we sorted each electrode separately. The spikes were sorted in the raw 76 dimensional space without dimensional reduction. For each sorted unit, the spike sorting algorithm had two phases. First, the algorithm estimated a suitable seed spike. Second, the corresponding waveform was optimized iteratively until the spike assignments of that unit remained constant. The clustering algorithm selected a seed spike by calculating the average noise level across all units. Afterwards, it randomly chose one spike and counted the number of neighboring spikes within this average noise level. Those spikes were called the spike-neighborhood. This procedure was repeated for 500 randomly chosen spikes in order to maximize the chance of finding a globally optimal seed spike. The spike that had most neighbors was selected as the seed for a unit. In order to optimize this spike seed, the noise level for the neighboring spikes was recalculated, the new neighborhood was calculated given this new noise level, and the new average waveform was calculated. This procedure was repeated until the neighborhood remained constant. The spikes within the noise-defined neighborhood were considered to belong to one sorted unit. For this unit, the spike sorting was finished at this point and it was not considered for further spike sorting. For the remaining spikes, the algorithm re-started phase one and two in order to search the next sorted unit. This procedure was stopped when it resulted in sorted units with spike rates lower than 0.1 Hz.

We regarded a unit as a single unit when the number of spikes within an inter-spike interval of less than 2 ms corresponded to a smaller firing rate than the average firing rate of the unit. To define the degree of decorrelation across neurons, we used the Μ-rate^20^. The Μ-rate denotes the minimum spike rate in the spike-triggered spike average between two neurons (cross correlogram). The cross correlogram was calculated over a period of -10 to 10 s with a 10 ms binning. We did not calculate the Μ-rate from a neuron to itself since that would reflect intra-neuronal processing (adaptation and refractory period) rather than the decorrelation of the population. The Μ-rate corresponds to the average spike rate if the spikes of the two neurons occur independently of each other, and the Μ-rate would be 0 for the case of a lag with no corresponding spike pairs. The Μ-rate percentage was calculated by dividing the Μ-rate with the average firing rate.

### Single and multiunit velocity modulation

As a general way to relate behavior to neural activity on a single unit or multiunit level, we used a generalized form of spike triggered average of the paw velocity, which we denote as activity weighted distribution (AWD). First, instead of taking discrete spikes, we weighted the behavioral variable (paw velocity or position) with a continuous neuronal activity. Here this continuous activity was the instantaneous firing rate smoothed with a Gaussian kernel with a standard deviation of 50 ms. Second, instead of averaging the behavioral variable, we calculated the distribution for the behavioral variable. A distribution was formed by binning the complete velocity range into 10 equally sized bins. Each bin quantified the average activity across the velocity range of the corresponding bin. In contrast to the linear average in the classical spike triggered average, the distribution of the behavioral variable allowed us to take nonlinearities into account, e.g. exponentially increasing firing rates with linearly increasing velocity. According to a traditional spike-triggered average, the relation between neuronal activity and behavior was calculated at different temporal lags between neural activity and behavior. Here we used lags between -4 and 4 s with a temporal resolution of 10 ms. For large delays beyond 3 s, the neuron was typically no longer modulated by behavior. Here we used the average activity between 3 and 4 s to calculate a baseline activity. This baseline activity was subtracted from the AWD. The average velocity modulation at each lag was calculated by taking the mean of the absolute value of the subtracted AWD (**Fig. 1F and G**). The duration and the lag of the modulation was calculated by first extracting the peak modulation. Then we traced this modulation backward and forward in time until the modulation was less than 80% of the peak modulation. The temporal difference between those two time points was defined as the duration of the modulation (**Fig. 1H, 1I, 2B, 2C, and 2D**). The average between those time points was denoted as the temporal lag of the modulation. We took the average time of the 80% start and stop time since this resulted in a more accurate estimation than the peak time. This was due to the frequent occurrence of plateaus in the velocity modulation. During these plateaus, small fluctuation of the neuronal signal within the noise level can make the peak appear at any time point along the plateau. To determine if a unit was modulated by velocity, we calculated the mean and standard deviation of the velocity modulation at the two extreme lags of the normalized velocity modulation (−4 to -3 s and 3 to 4 s). The normalized velocity modulation was calculated by subtracting and dividing the velocity modulation with the mean and standard deviation, respectively. A unit was regarded as modulated if this velocity modulation was larger than 10 (a.u.).

### Bootstrapping velocity modulation

To estimate the variability of the modulation duration we used a bootstrap analysis (**Fig. 1H and I**). Since it would be computationally inefficient to sample from all 10 ms bins with replacement and since 2 neighboring 10 ms bins were not independent, we chose to divide each session into 100 segments of equal size and to calculate the AWD for each such segment. This resulted in segments that were at least 10 seconds long, allowing computationally effective bootstrap sampling. We sampled the corresponding 100 AWDs with replacement and calculated the resulting velocity modulation. This procedure was repeated 100 times. For each repetition, we calculated the modulation duration. Afterwards, we calculated the standard deviation across those repetitions.

### Population correlation analysis and trial definition

The population correlation analysis was performed on normalized neural activity. For each unit, we divided the spike trains into 10 ms bins, subtracted the average firing rate and divided each bin by the standard deviation of the binned activity. This normalized data was organized into a matrix with as many rows as there were units and as many columns as there were time bins. To prepare the data for the correlation, we normalized each column to have an average of 0 and a Cartesian norm of 1 (unit length). Finally, we removed a global population activity that could otherwise bias the correlation analysis. During short periods of time (between 500 ms to 10 s) sometimes the animals suddenly froze (both in the joystick and the locomotor task) which resulted in a correlated population activity across the joystick and the locomotor task (average R=0.5). Since this activity was correlated across two fundamentally different tasks, it was more likely to reflect a global state change rather than a planning process, which in turn could bias the population correlation. Therefore, we minimized the contribution of this freezing related population activity, p, by correlating the population activity at each time bin, a_t_, with the population activity, and subtracting the population activity according to this correlation: a_t_ – p(a_t_*p), where * is the scalar product.

With this normalized activity, we calculated the scalar product (Pearson correlation coefficient) between two population vectors at 2 different time points (**Fig. 3C and D**). We only correlated population vectors within a trial. Since our behavioral data was not separated into defined trials, we constructed trials using the paw velocity. First, we filtered the paw velocity with a Gaussian kernel of 2 s full width half maximum (FWHM). To find trials for which a period of low behavioral activity was followed by a period of high behavioral activity, we divided each time point in the filtered velocity by each time point in the filtered velocity 2 s earlier. If this ratio was larger than 2 and if this ratio was a local maximum across time, this was regarded as the central time point of a trial. A trial was then defined as 8 s before and 8 s after this maximum. This resulted in 1601 bins of 10 ms in one trial. The correlation was calculated between all 1601×1601 pairs of time points within a trial. Finally, as the population vector at one reference time point was correlated with the population vector at all other time points, the correlation would decay with increasing distances from the reference time point. This decay was fitted by an exponential function using nonlinear optimization with a Gaussian cost function (**Fig. 3E, F, G and H**).

### Behavioral impact on population correlation

To test how well the neurons encoded for position (**Fig. S2B**), we divided the egocentric x and y movement coordinates of the right paw into five equally sized bins between the minimum and maximum position value. This resulted in a 5 × 5 element matrix. For each element in this matrix we calculated the average firing rate of the neuron when the paw was in the corresponding position within ±50 ms. We used this matrix as a lookup table to estimate the instantaneous firing rate at each 100 ms time bin, given the position at the corresponding time bin. The resulting time course of the firing rate was correlated to the time course of the true instantaneous firing rate binned in 100 ms bins. The same analysis sequence was conducted for x and y velocity.

### Subthreshold reconstruction

The subthreshold reconstruction algorithm, SubLab, has been described in detail elsewhere^20^. In short, the algorithm uses the spikes of one unit (target unit) to reconstruct its subthreshold activity by means of the spiking activity of the remaining units (input units). The algorithm differs from recent auto-encoders and dimension reduction techniques in three aspects: (1) it does not assume an even distribution of spikes in time (Poissonian or Gaussian models); (2) (subthreshold) activity is not modified, as long as it does not cross the threshold; (3) the algorithm reconstructs the subthreshold activity individually per neuron and, therefore, does not impose any relation between units. Here we used 10 training epochs and we ran the reconstruction on complete sessions.

We also tested the LFADS auto-encoder algorithm, since it does not require a trial structure and since it can fit complex dynamics to spiking data. For our data, LFADS smoothed the spike trains in a piecewise continuous way. We observed gaps in the smoothed spike trains. We suspect that these gaps were due to the spontaneous and complex behaviors, which in turn caused the internal states to be reset frequently.

The reconstructed activity was filtered in the following way (**Fig. 4C, D, E and F**). High pass filtering: First, the reconstructed signal was smoothed with a Gaussian kernel with a standard deviation (σ) of 0.14 s. Using the cut-off frequency formula for Gaussian filtering (2πs)^-1^, this corresponds to a cut off frequency of 1.1 Hz. Second, we subtracted this smoothed signal from the original reconstructed signal. Band-pass filtering: First, the reconstructed signal was smoothed with a Gaussian kernel with a standard deviation of 0.057, 0.14, 0.28, 0.57, 1.4, 2.8, and 5.7 s (2.8, 1.1, 0.57, 0.28, 0.057, and 0.028 Hz), respectively. Second, we subtracted this smoothed signal from the original reconstructed signal. Third, the resulting signal was smoothed with a Gaussian kernel with a standard deviation of 0.014, 0.035, 0.071, 0.14, 0.35, 0.71, and 1.4 s (11, 4.5, 2.2, 1.1, 0.45, 0.22, and 0.11 Hz), respectively. Low pass filtering: The band-pass filtered signal that was filtered with a low-pass kernel of 0.71 seconds (0.22 Hz) and high-pass kernel of 2.8 seconds (0.057 Hz) was referred to as the low-pass filtered signal.

The additional high pass filtering minimizes the influence from strong low frequency components. Finally, to get the energy of the filtered signal, we calculated the absolute value of the high-pass filtered signal.

### Relating population and frequency coding

Output-null and output-potent coding has traditionally been studied during the planning and execution phase of an instructed delay tasks. Since our behavioral setting does not include a typical trial structure, we defined the planning and execution phase in terms of the lag between the paw velocity and the neuronal activity. To this end, the anterior-posterior paw velocity was multiplied with the neuronal activity for a sorted unit in a bin-wise manner for a given lag and averaged across all bins. Thus for a given lag, this approach will quantify how each neuron codes for the movement in a linear manner. We used temporal lags from -1 second to 1 second with 10 ms bins. The result is a N x 201 dimensional matrix for each session, where N is the number of sorted units. Dimension reduction to a 2×201 matrix was achieved by taking the largest two principal components. We defined the output potent space as a one-dimensional space covered by the vector between origo (0, 0) and the point in the two dimensional space (spanned by the first two principal components) at lag of -40 ms. This lag was defined by the notion that executional activity should have a small correlation to the planning activity, which in turn translates to the lag at which the correlation to the average activity between - 1000 and -200 ms (putative planning activity) was smallest. We choose the upper limit to be -200 ms to minimize the bleeding into executional activity^8,9^. Since this definition of the lag for the output potent space is maximizing the separation of planning and executional activity it is maximizing the chance that the null and potent spaces will be found. Such a biased definition is justified here since the aim is not verify the null space theory but rather to see if it is related to the ratio of high and low frequencies of neuronal changes. The output null space was orthogonal to this output potent space. For each lag, we estimated which state the neuronal activity was closest to by means of the difference in magnitude: abs(output potent)-abs(output null). If this ***<u>state tuning</u>*** was positive we regarded the neuronal activity to be in the output potent state and if it was negative we regarded the neuronal activity to be in the null space.

To test if the frequency coding can predict whether the neuronal activity is in the output potent or output null space, we assigned the frequency preference for each lag of a certain session. This was done by calculating the difference in magnitude: abs(amplitude of high frequency) – abs(amplitude of low frequency). If this ***<u>frequency tuning</u>*** was positive, it means that the neuronal activity had a larger high frequency component and if it is negative it means that the neuronal activity had a larger low frequency component. We pooled all lags (across all sessions) that had a positive frequency (or negative frequency) tuning and calculated the resulting average state tuning.

### Behavioral quantification during optogenetic stimulation

For optogenetic stimulation we used a 200 Μm fiber implanted at 1 mm depth in the primary motor cortex of two rats (511 and 512) (AP=0.5, LM=2, and DV=1). The viral vector AAV5 carrying the construct hSyn-hChR2(H134R)-eYFP-WPREpA (UNC vector core, Chapel Hill, NC, USA), was injected at a depth of 1.5 mm with a volume of 1Μl. Each stimulation trial lasted 10 seconds and the light intensity was sinusoidally modulated according to one of five frequencies: 0.1, 0.3, 1, 3, and 10 Hz with a peak power of 4-12mW at the fiber tip. Since the current that ChR2 can give rise to is smaller for low frequencies than for high frequencies, we compensated with a stronger light intensity for the lower frequencies^32^. Each trial was randomly interleaved with 120 to 240 seconds.

To quantify the subtle paw movements that results from sinusoidal optogenetic stimulation (Fig. 6C), we first calculated the paw position using FreiPose^31^. For a given trial we manually selected the camera with the clearest view of the right foot (the optogenetic stimulation was in the left hemisphere). The paw position for each frame was then projected to this camera and a 100×100 pixels window was cut out around this projected position. The optical flow was calculated for each pair of neighboring frames (opticalFlowHS object in Matlab). The paw position for both frames in this pair was taken according to the first frame. The vertical component of the optical flow was extracted since this is the major movement axis during stimulation. Finally, the optical flow was only sampled at pixels with a saturation above 20% (0.2 for saturation of the rgb2hsv function in Matlab). This was done in order to sample paw movements rather than more unspecific “fur” movements. Trials in which the rat was grooming, eating or walking was eliminated from further analysis. The amount of movement for each stimulation frequency was then calculated by averaging the energy in the 0.1, 0.3, 1, 3 and 10Hz band (using the spectrogram function in Matlab with window size of 100 and overlap 99).

In addition to the automatic behavioral quantification described in the previous paragraph, in figure 6D we manually quantified how the animal responded to the optogenetic stimulation. To this end we measured the duration for which the rat performed an abnormal behavior. Abnormal behavior was defined as a cyclic paw movement for which the rat was lifting and lowering the right paw at least one time. We excluded movements that could be ascribed to walking, grooming or movements that showed a coordination between left and right paw. Although the cyclic criterion seemed robust, there was one trial in which rat 512 stretched out the paw abnormally for the 10 Hz stimulation and this was therefore not counted as a cyclic movement.

### Statistical procedures

All statistics and graphical illustrations of spiking unit data have been corrected for the possibility that the same unit has been recorded during multiple consecutive days (**Supplementary Table 3**). In motor cortex, evidence has been provided that tungsten electrodes are able to record the same unit for an average of three days^33^. Since a considerable amount (11%) of neurons could be recorded for up to a week, we regarded every 7^th^ unit to be an independent data sample. To this end, the degrees of freedom were calculated on the basis of the unit count divided by 7. We made this correction for the t-test, the Pearson correlation coefficient, and the ANOVA. For box plots (using Matlab’s boxplot function), we plotted the bootstrapped data (using Matlab’s bootstrap function with 1000 iterations) and adjusted the standard deviation of the bootstrapped data such that it was 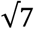 times that of the original data.

For statistical testing, we assumed that the data was normally distributed. The test statistics for the Pearson correlation coefficient, the ANOVA and unpaired statistics approached a normal distribution for large data samples. For the paired t-test, we assumed a normal distribution as the test distribution was symmetric around 0. Unless otherwise stated, samples were described as mean and standard deviation of the mean.

Since we had one less animal in the joystick task (animal 220 lost the implant before it learned the joystick task), all paired tests were done without animal 220 in both the joystick and locomotor task. The non-paired tests were done using all 6 animals in the locomotor task and all 5 animals in the joystick task.

## Acknowledgements

We would like to thank Philippe Coulon for comments on earlier versions of the manuscript. This work was supported by the Bernstein Award 2012 (01GQ2301), the cluster of excellence BrainLinks-Brain-Tools (EXC 1086), the Deutsche Forschungsgemeinschaft (DFG) via the grants DI 1908/3-1 and 1908/6-1, as well as the ERC Starting Grant OptoMotorPath (338041), all to I.D. The authors acknowledge support by the state of Baden-Württemberg through bwHPC and the German Research Foundation (DFG) through grant no INST 39/963-1 FUGG (bwForCluster NEMO).

## Author contributions

D.E. and I.D. conceived and designed the experiments and wrote the manuscript. D.E., M.H., and A.S. performed the experiments.

## Data Availability

The data that support the findings of this study are available from the corresponding authors upon reasonable request.

## Additional Information

Supplementary Information is available for this paper.

Correspondence and requests for materials should be addressed to.

