## Supplementary Infomration for "Cortical activity at different time scales: high-pass filtering separates motor planning and execution"

**Supplementary Information**

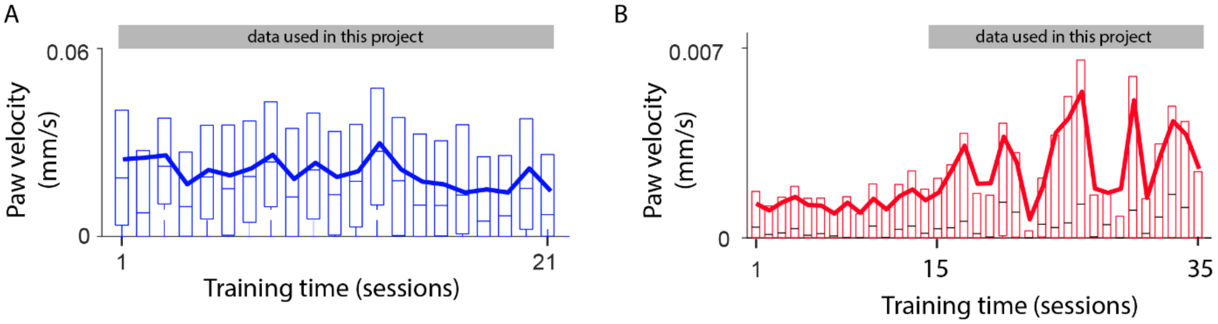


**Supplementary Figure 1.** Development of the paw velocities across recording sessions. A: Paw velocity for the locomotor task. Outliers and upper whisker artefacts have been removed from the box-plot to improve visibility. The solid line is the average paw velocity. The gray bar on the top indicates which data has been used for this project. B: Same outline as in A but for the joystick task. The paw velocity changed at session 15. Thus, data was used only after 15 training sessions.

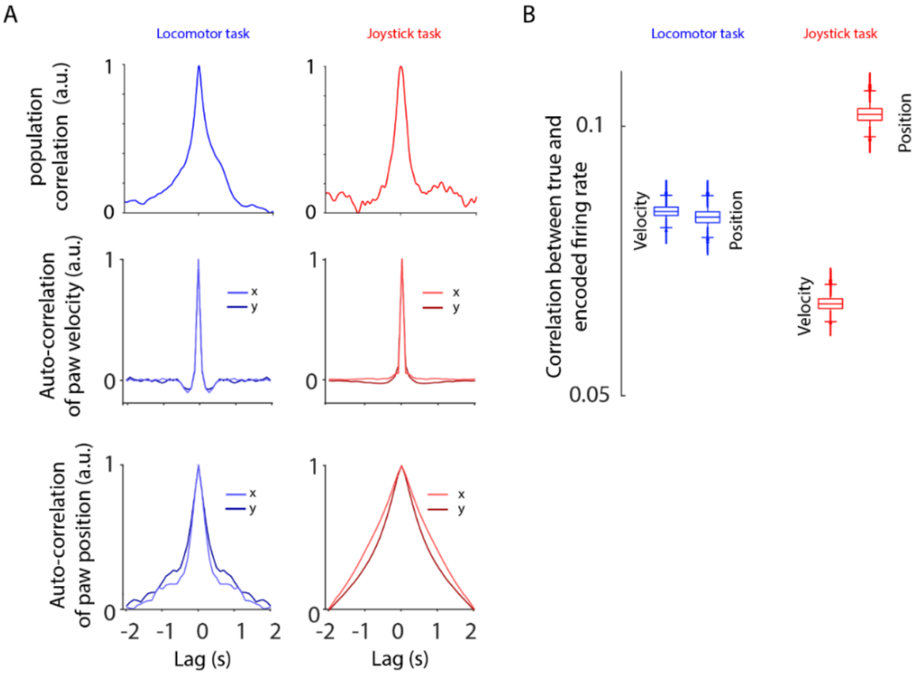


**Supplementary Figure 2.** Ruling out a putative behavioral impact on population correlations. A: Population activity correlation for the locomotor task and the joystick task (top row). Autocorrelation of the paw velocity for the locomotor task and the joystick task (second row), and of the paw position for the locomotor task and the joystick task (bottom row). B: Encoding performance using only position or direction of the right paw.

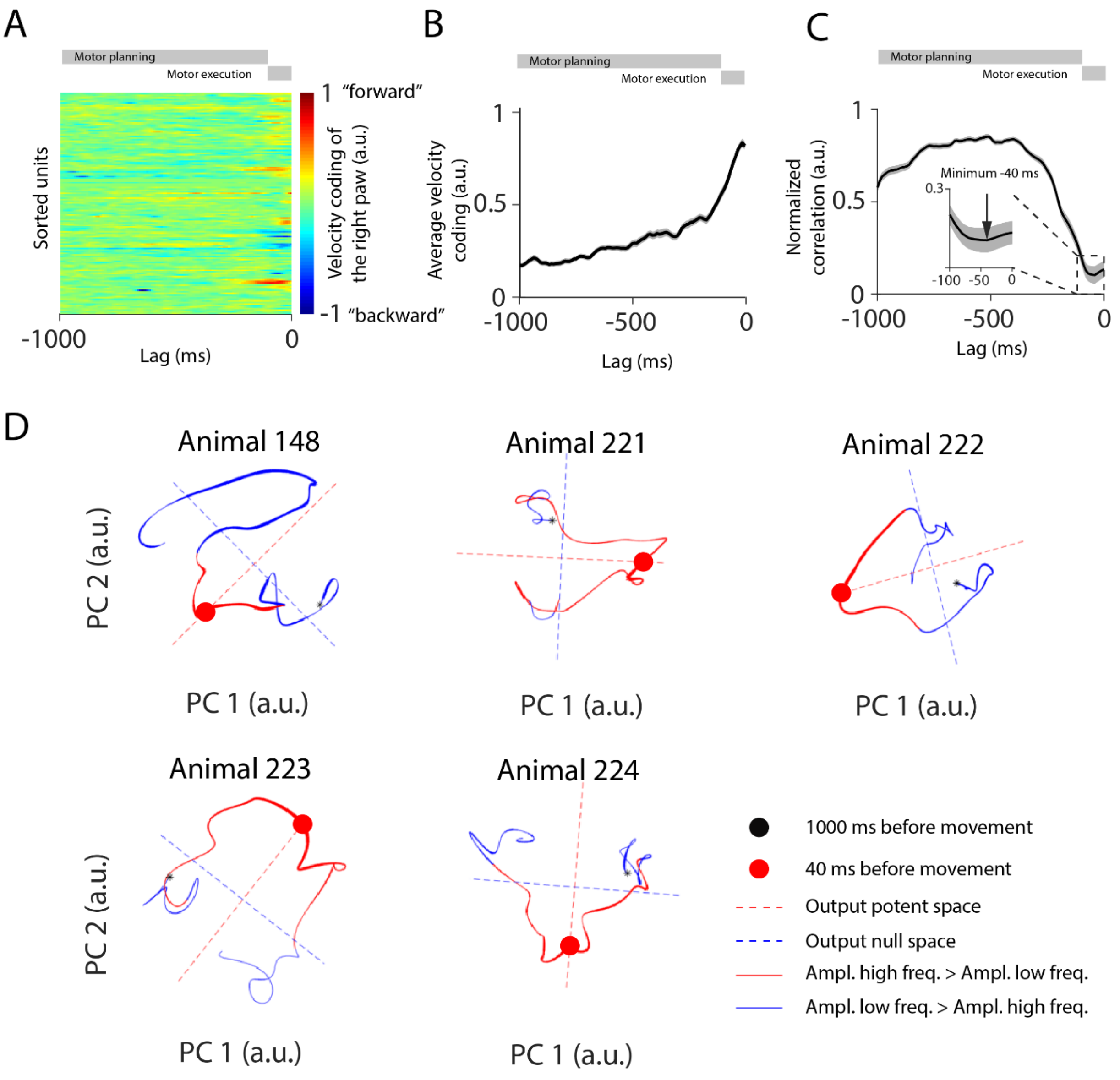


**Supplementary Figure 3**. Ratio of high to low frequency changes of neuronal data predict state spaces. A: Velocity tuning across all units in one session of rat 222 across temporal lags ranging from -1000 to 1000 milliseconds. B: Average velocity tuning across all units, all sessions of all animals. C: Correlation between the average population activity between -1000 and -200 ms and the population activity at all lags between -1000 and 0 ms (x-axis). The correlation reaches a minimum at -40 ms (inset). This minimum describes when the motor execution related activity is maximally different from motor planning related activity and is hence used to define the output potent space. D: Example sessions of dimension reduced population coding for the five animals in the joystick task. The trajectory is divided into paths for which the high frequency (>1.1Hz) had a larger amplitude than the low frequency (<1.1Hz) (red), and into segments for which the high frequency had a smaller amplitude than the low frequency (blue). Output null (blue) and output potent spaces (red) are indicated by dashed lines. The thickness of the trajectory indicates the averaged velocity tuning (B). Thicker lines refer to stronger correlation between paw velocity and neuronal activity. All animals apart rat 148 showed a clear correlation of the output potent space with paw velocities. This rat had the lowest signal to noise ratio and the electrodes covered a smaller cortical area.

**Supplementary Table 1**

| Animal | SU mod | SU all | MU mod | MU all | SU+MU mod | SU+Mu all | Session count |
|---|---|---|---|---|---|---|---|
| 148 | 162 | 565 | 463 | 852 | 625 | 1417 | 15 |
| 220 | 199 | 493 | 410 | 706 | 609 | 1199 | 11 |
| 221 | 86 | 519 | 337 | 858 | 423 | 1377 | 16 |
| 222 | 367 | 1321 | 679 | 1566 | 1046 | 2887 | 21 |
| 223 | 706 | 1816 | 893 | 1708 | 1599 | 3524 | 20 |
| 224 | 212 | 686 | 589 | 1186 | 801 | 1872 | 17 |
| Sum | 1733 | 5400 | 3371 | 6876 | 5103 | 12276 | 100 |

Counts of neuronal units recorded during the locomotor task. mod - modulated units.

**Supplementary Table 2**

| Animal | SU mod | SU all | MU mod | MU all | SU+MU mod | SU+MU all | Session count |
|---|---|---|---|---|---|---|---|
| 148 | 30 | 217 | 73 | 266 | 103 | 483 | 5 |
| 220 | 0 | 0 | 0 | 0 | 0 | 0 | 0 |
| 221 | 9 | 161 | 53 | 253 | 62 | 414 | 5 |
| 222 | 72 | 344 | 238 | 536 | 310 | 880 | 7 |
| 223 | 63 | 324 | 175 | 349 | 238 | 673 | 4 |

| | | | | | | | |
|---|---|---|---|---|---|---|---|
| 224 | 28 | 171 | 98 | 255 | 126 | 426 | 4 |
| Sum | 202 | 1217 | 637 | 1659 | 839 | 2876 | 25 |

Counts of neuronal units recorded during the joystick task. mod - modulated units.

### Supplementary Table 3

| Fig. | Groups:<br>**a** refers to animal count,<br>**u** refers to sorted unit count,<br>**$u_c$** = u/7 refers to corrected unit count used for significance | Statistical Analysis |
|---|---|---|
| Fig. 1l | Locomotor task, Motor-Sensory, (u = 5103, $u_c$=729)<br>Joystick task, Motor-Sensory, (u = 839, $u_c$ = 120) | Two-tailed paired t-test, $P < 0.0001$<br>Two-tailed paired t-test, $P < 0.0001$ |
| Fig. 2d | Locomotor task, Absolute lag, (u = 5137, $u_c$=729)<br>Locomotor task, Duration (6 bins), (u = 5137, $u_c$=729) | RM One-Way ANOVA, $F\ (5, 723) = 474$, $P < 0.0001$ |
| Fig. 2d | Joystick task, Absolute lag, (u = 839, $u_c$ = 120)<br>Joystick task, Duration (6 bins), (u = 839, $u_c$ = 120) | RM One-Way ANOVA, $F\ (5, 114) = 72$, $P < 0.0001$ |
| Fig. 2d (left) | Locomotor task, S1 Duration, (u = 1651, $u_c$=236)<br>Locomotor task, M1 Duration, (u = 2316, $u_c$=331)<br>Locomotor task, M2 Duration, (u=1136, $u_c$=163) | RM One-Way ANOVA, $F\ (2, 726) = 171$, $P < 0.0001$<br>Post Hoc: 3-way Bonferroni:<br>S1-M1: p=0.0021<br>M1-M2: p=0.024<br>S1-M2: p<0.0001 |
| Fig. 2d (right) | Joystick task, S1 Duration, (u = 252, $u_c$=36)<br>Joystick task, M1 Duration, (u = 323, $u_c$=46)<br>Joystick task, M2 Duration, (u= 264, $u_c$=38) | RM One-Way ANOVA $F\ (2, 117) = 16$, $P < 0.0001$<br>Post Hoc: 3 way Bonferroni:<br>S1-M1: p<0.0001<br>M1-M2: p<0.0001<br>S1-M2: p<0.0001 |
| Fig. 3f | Locomotor task, Relative time constant, (a = 6)<br>Joystick task, Relative time constant (a = 5) | Two-tailed t-test, $P = 0.20$<br>Two-tailed t-test, $P = 0.043$ |
| Fig. 3h | Locomotor task, Absolute time constant, (a = 6)<br>Joystick task, Absolute time constant (a = 5) | RM One-Way ANOVA, $F\ (3, 18) = 6.6$, $P = 0.0033$<br>Post Hoc: 6-way Bonferroni:<br>Lowest paw velocity in the Joystick task vs Highest paw velocity in the Locomotor task: p = 0.037<br>Lowest paw velocity in the Joystick task vs Highest paw velocity in the Joystick task: p = 0.004 |
| Fig. 4d | Locomotor task, low-pass versus high-pass, Pearson correlation, (a = 6)<br>Joystick task, low-pass versus high-pass, Pearson correlation, (a = 5) | Two-tailed t-test, $P = 0.016$<br>Two-tailed t-test, $P = 0.0091$ |
| Fig. 4d | Locomotor task, high-pass, Lag, (a = 6)<br>Joystick task, high-pass, Lag, (a = 5) | Two-tailed t-test, $P = 0.33$<br>Two-tailed t-test, $P = 0.01$ |
| Fig. 4d | Locomotor task, low-pass, Lag, (a = 6)<br>Joystick task, low-pass, Lag, (a = 5) | Two-tailed t-test, $P = 0.0013$<br>Two-tailed t-test, $P = 0.0029$ |
| Fig. 4e | Locomotor task, Pearson correlation, (a = 6) | RM One-Way ANOVA $F\ (6, 35) = 3.6$, $P = 0.007$<br>Post Hoc: 21-way Bonferroni:<br>5 Hz vs 1 Hz: p = 0.04 |
| Fig. 4e | Joystick task, Pearson correlation, (a = 6) | RM One-Way ANOVA $F\ (6, 28) = 3.5$, $P = 0.01$ |

| | | |
|---|---|---|
| | | Post Hoc: 21-way Bonferroni:<br>10 Hz vs 0.5 Hz: p = 0.034<br>5 Hz vs 0.5 Hz: p = 0.034 |
| Fig. 4f | Locomotor task, Lag, (a = 6) | RM One-Way ANOVA F (6, 35) = 5.8, *P* = 0.0002<br>Post Hoc: 21-way Bonferroni:<br>50 Hz vs 0.5 Hz: p = 0.0021<br>20 Hz vs 0.5 Hz: p = 0.042<br>10 Hz vs 0.5 Hz: p = 0.0017<br>5 Hz vs 0.5 Hz: p = 0.0089<br>2 Hz vs 0.5 Hz: p = 0.0009 |
| Fig. 4f | Joystick task, Lag, (a = 6) | RM One-Way ANOVA F (6, 28) = 2.5, *P* = 0.045<br>Post Hoc: 21-way Bonferroni:<br>10 Hz vs 0.5 Hz: p = 0.045 |

**Supplementary Note 1:** *The difference in stability of population activity cannot be explained by behavioral differences across the two tasks*

Could the differences in stability of the population activity be explained by differences in behavior across the two tasks? To address this question, we tested whether the autocorrelation of two easily accessible behavioral parameters can explain the observed effects: (1) paw velocity and (2) egocentric paw position. As the population correlation decayed more slowly in the locomotor task, we would expect a temporally broader behavioral autocorrelation for the locomotor task compared to the joystick task. However, the similarly narrow peaks of the autocorrelation of the paw velocity in both tasks argue that the difference in the stability of the population activity (**Supplementary Fig. 3A, upper panel row**) cannot be explained by differences in paw velocity (**Supplementary Fig. 3A, middle panel row**). Similarly, for the paw position, we would expect a temporally broader behavioral autocorrelation for the locomotor task compared to the joystick task to explain the stability differences of the population activity. Instead, the autocorrelation of the paw position was narrower during the locomotor task than during the joystick task (**Supplementary Fig. 3A, lower panel row**). Thus, neither the velocity autocorrelation, nor the position autocorrelation, could explain the differences in population activity stability. Alternatively, neurons preferentially encode the position during the locomotor task, and the velocity during the joystick task. If this were true, the broad population correlation during locomotion could be explained by the broad position autocorrelation and the narrow population correlation during the joystick task could be explained by the narrow velocity autocorrelation. To this end we tested the encoding preference (position or velocity) of the neurons in the two tasks. The neurons showed an encoding preference for position in the joystick task (**Supplementary Fig. 3B**), which stands in contrast to the more precise auto-correlation for paw velocities in the joystick task. These opposing results suggest that differences in the stability of population activity cannot be explained by a differential encoding preference of position and velocity. To summarize, there is a strong decorrelation during paw movements in the joystick task, which cannot be explained by means of differences in behavioral statistics.

**Supplementary Note 2:** *Adaptation mechanisms and compatibility with prolonged movements*

Adaptation mechanisms at any stage between the cortex and the muscles could serve as the biological equivalent of a high pass filter. The high pass filter should detect fast changes in the activity. On the level of neuronal spiking this can be a change from a high firing rate to a low firing rate, or vice versa. On the level of summed synaptic input this can be the change from a large input current or a low input current, or vice versa. There are numerous neuronal phenomena that describe high-pass filtering on the time scale of hundreds of milliseconds, such as spike rate adaptation[1], short term synaptic depression[2], integrating inhibitory neurons[3], low-pass filtering across gap junction connected interneurons[4], and depolarization block[5]. The underlying mechanisms of those phenomena can follow the slowly evolving motor planning and sensory integration activity by means of the intracellular calcium concentration, amount of release ready vesicles in the presynaptic terminal, the firing rate of integrating inhibitory neurons, or the number of inactivated sodium channels, respectively.

The generation of prolonged movements under the control of an adaptation related high pass mechanism would require a subcortical process that can be triggered by short lasting inputs. In the lamprey, the reticulospinal cells transform a short duration sensory input into a long-lasting excitatory command[6]. In the zebrafish, high frequency stimulation in the brain stem initiates sustained locomotor behavior[7]. Similarly, in the basal ganglia of rodents, neurons are activated during the initiation and termination of movement sequences[8]. In the mouse, the lower pyramidal tract neurons have been shown to have a preference for motor execution[9] and are thus good candidates for contributing to sustaining movements. These neuronal processes in combination with our proposed high pass filtering mechanisms would allow for movements of different durations.
